# Structural insights into the distinct protective mechanisms of human antibodies targeting ZIKV NS1

**DOI:** 10.1101/2023.10.16.562450

**Authors:** Qi Pan, Xiaomin Xing, Jianhai Yu, Qiang Chen, Haizhan Jiao, Wanqin Zhang, Yingfen Wen, Ming Gao, Wei Zhao, Lei Yu, Hongli Hu

**Affiliations:** Kobilka Institute of Innovative Drug Discovery, School of Medicine, The Chinese University of Hong Kong, Shenzhen, Guangdong, 518172, China; Guangzhou Eighth People’s Hospital, Guangzhou Medical University, Guangzhou, Guangdong, 510060, China; BSL-3 Laboratory (Guangdong), Guangdong Provincial Key Laboratory of Tropical Disease Research, School of Public Health, Southern Medical University, Guangzhou, Guangdong, 510515, China

## Abstract

Zika virus (ZIKV) Non-structural protein 1 (NS1) plays an essential role in viral replication and immune evasion. Our understanding of the differential protective mechanism of NS1-targeting antibodies is limited. Here, we determined the cryoEM structures of ZIKV NS1 in complex with two group antibodies at 2.6-2.9Å. Group I antibodies (3G2 and 4B8) potently recognize cell surface form of NS1 and multiple oligomeric forms of NS1 by occupy the epitopes on outer surface of dimeric NS1. IgG and Fab from group I antibodies completely abrogate sNS1-mediated endothelial dysfunction in vitro. Group II antibodies (4F10, 2E11, and 14G5) recognize the previously reported epitopes in distal end of the *β*-ladder domain of monomeric NS1, and their blockade efficiency depends on the affinity with NS1 protein. These findings elucidate the correlation between the epitope recognition and the protective efficacy of anti-NS1 antibodies and highlight the distinct mechanisms of therapeutic potential of 3G2 and 4B8.

## Introduction

Zika virus (ZIKV) is a mosquito-borne pathogen that belongs to the flavivirus genus of Flaviviridae, and other flavivirus belonging to this family include Dengue Fever virus (DENV), West Nile virus (WNV), Yellow Fever virus (YFV) and Japanese encephalitis virus (JEV)^1^. Since its first identification in Uganda in 1947, ZIKV has caused several global outbreaks and has become a significant health concern^2^. During pregnancy, ZIKV infection cause infants being born with microcephaly and other congenital malformations. In adults, it has been linked to Guillain-Barré syndrome, neuropathy, and myelitis^3^. Unfortunately, there is no approved vaccine or therapeutic against ZIKV^4^.

Non-structural protein 1 (NS1) is gaining attention as a promising therapeutic target^5,6^. The NS1 gene in flaviviruses is highly conserved and encodes a 352 amino acid protein with three distinct domains: an N-terminal *β*-roll domain (residues 1-29), a wing domain (residues 30-180), and a *β*-ladder domain (residues 181-352) ^7^. During infection, NS1 was initially found as a monomer in plasma, subsequently associated with the intracellular membrane and the cell membrane surface as a dimer, and eventually secreted into the bloodstream as a hexamer^8,9^. Soluble secreted NS1 (sNS1) is an important diagnostic marker for viral infection as it circulates in high concentrations in virus-infected patients. As a multifunctional factor, NS1 can facilitate viral RNA replication and virus assembly, inhibit complement^10^, activate platelets and immune cells^11,12^, and induce vascular endothelial dysfunction in a tissue-specific manner^13^. ZIKV sNS1 may also promote placental dysfunction via modulation of glycosaminoglycans on trophoblasts and chorionic villi^14^. Several studies have showed that NS1-based vaccines are capable of inducing robust humoral and cellular immunity in mice^5,15-18^. Passive transfer of NS1 antibodies provides in vivo protection via activating Fc-mediated effector functions without inducing ADE effect^19,20^. Moreover, emerging evidence indicates that mAbs targeting ZIKV NS1 provide protection independent of Fc effector function^6,21^. Numerous epitopes of protective antibodies against NS1 have been mapped, showing the most antigenic regions are located in the wing domain and the C-terminal tip of the *β*-ladder^22^. Structural studies are focused on protective antibodies that targeting the *β*-ladder domain^23-25^. In addition, antibodies targeting epitopes on the cell-surface form of NS1 are thought to confer protection against ZIKV infection^20,26^.

However, our understanding of the protective mechanisms of antibodies that recognize distinct epitopes is restricted due to a deficiency in structural details.

In a previous study, Gao et al. isolated a series of NS1-specific mAbs (3G2, 4B8, 4F10, 2E11, and 14G5) from ZIKV convalescent patients^27^. Among these mAbs, the protective efficiency of 4F10 (targeting the Cterminal region) exhibited weaker protective efficacy that relied on Fc-mediated effector function, while 3G2 and 4B8 (targeting the N-terminal region) provided superior protection via both the Fc-dependent and the Fc-independent pathways^6^. Here, we determined high-resolution structures of full-length ZIKV NS1 combined with five NS1-targeted human antibodies (tNS1-3G2, tNS1-4B8, tNS1-4F10, dNS1-2E11, and dNS1-14G5). Based on the distinct binding sites, we categorized them them into two groups. Combining with biochemical analysis, we elucidate the correlation between the unique recognition mode and the protective efficacy of NS1-targeted antibodies. These data providing significant clues for rational design of NS1-based vaccines and therapeutics.

## Results

### Recombinant ZIKV sNS1 are Mainly Homo-tetramers

The baculovirus expression system was used to express the full-length ZIKV NS1 (residues 3-352, GZ02 strain). After purification of sNS1 from *Sf*9 cell supernatant by affinity chromatography, SDS-PAGE and size-exclusion chromatography (SEC) analysis showed the purified NS1 is homogeneous (Extended Data Fig. 1a,b). Subsequent negative staining EM analysis indicates that purified sNS1 forms oligomers instead of dimers (Extended Data Fig. 1c). The two-dimensional (2D) class averages and three-dimensional (3D) initial model reconstruction show that the purified sNS1 mainly exists in the form of tetramer (tNS1) (Extended Data Fig. 1d,e). The tNS1 sample was applied for the subsequent complex preparation and the cryo-electron microscopy analysis.

**Fig 1.**
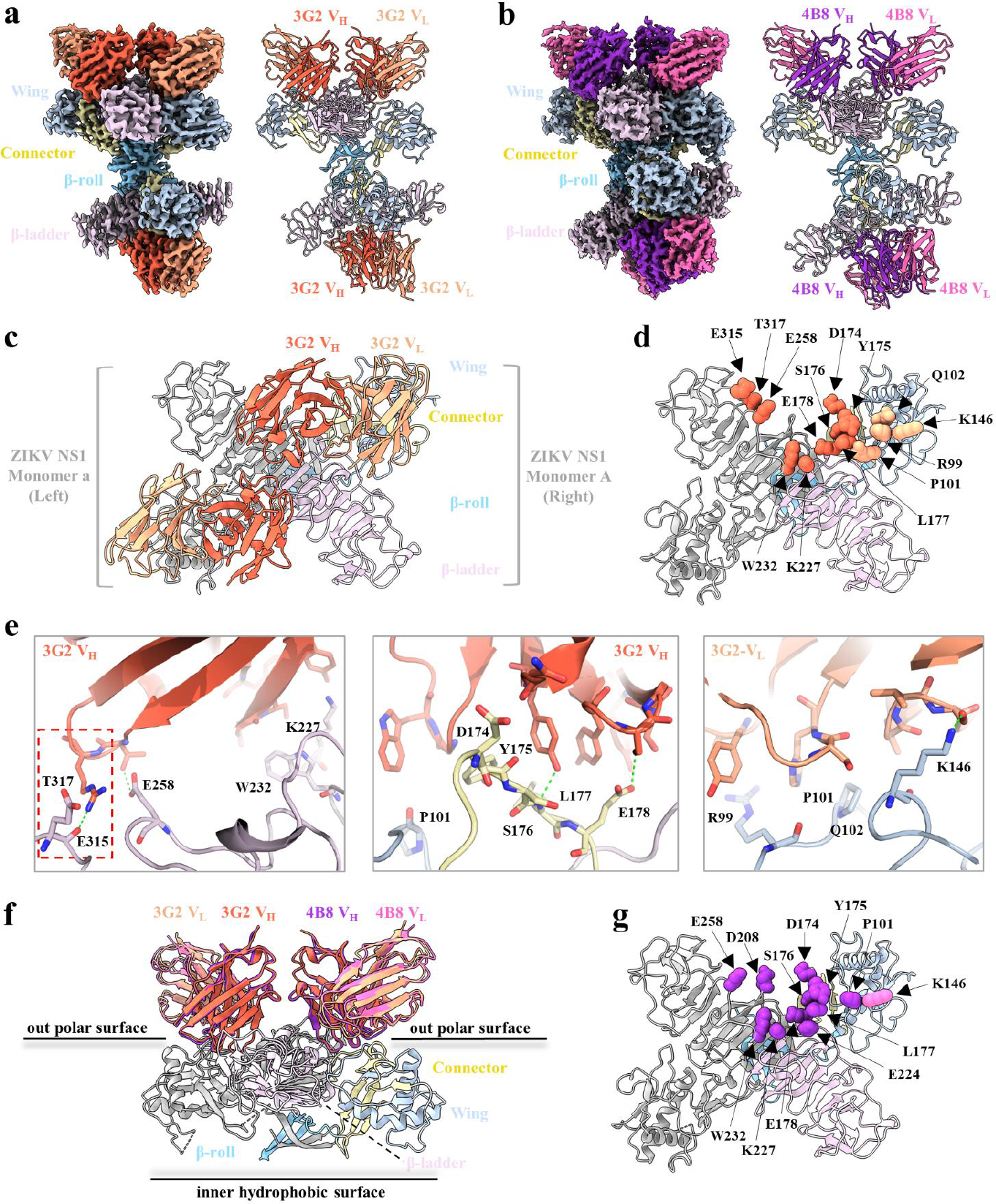
CryoEM structure analysis of ZIKV NS1 tetramer in complex with Fab 3G2 and Fab 4B8; (a, b) Overview of the density map and atomic model of the ZIKV NS1 tetramer with four Fab 3G2 variable region fragments and four Fab 4B8 variable region fragments; In each map and structure, the *β*-roll domain (residues1-29) of NS1 is colored light sky blue, connector (residues 30-37 and residues 152-180) is colored pale goldenrod, the wing domain (residues 38-151) is colored light steel blue and the *β*-ladder domain (residues 181-352) is colored thistle. Fab 3G2 is colored tomato (3G2 V_H_) and light salmon (3G2 V_L_). Fab 4B8 is colored dark orchid (4B8 V_H_) and hot pink (4B8 V_L_). V_H_, variable region of immunoglobulin heavy chain; V_L_, variable region of immunoglobulin light chain; This color scheme is used throughout the manuscript unless otherwise noted. (c) Zoomed-in view of the atomic model of the ZIKV NS1-3G2 complex; In this structure, the left monomer is dark gray, and the right monomer is color-coded by domain as in panel a. (d) Mapping of critical residues for 3G2 on ZIKV NS1 dimer in top view. The dimer is color-coded by domain as in panel c. Critical residues are indicated by spheres and are color-coded according to the mAb in panel c. The epitopes of 3G2 span two protomers of dimeric NS1. (e) Close-up view of the 3G2 epitope. Residues were shown as sticks; oxygen is colored red and nitrogen is colored blue and hydrogen bonding is represented by dotted green line. 3G2 V_H_ epitopes (left and middle), 3G2 V_L_ epitopes(right); The red box represents the distinct epitopes occupied by 3G2 V_H_ compared to 4B8 V_H_. This panel was generated using PyMOL software. (f) Comparison of binding modes for 3G2 and 4B8. Both 3G2 and 4B8 interact with the out polar surface of dimeric NS1, and they do not generate steric hindrance at the interface between the inner hydrophobic surface of the dimeric NS1 and the membrane. (g) Mapping of critical residues for 4B8 on ZIKV NS1 dimer in top view. The dimer is color-coded by domain following panel c. Critical residues are indicated by spheres and are color-coded following to mAb panel in B. The epitopes of 4B8 span two protomers of dimeric NS1. Except for panel e, all panels were prepared using ChimeraX.

### Group I antibodies (3G2 and 4B8) Target Epitopes Residues Span the Outer Surface of Dimeric NS1

To elucidate the NS1 antigenic sites and how anti-NS1 antibodies protect against ZIKV infection, we incubate the tetrameric NS1 with the antibodies 3G2, 4B8, 4F10, 2E11, and 14G5, and then further purified the complexes for structure determination by cryo-EM (Extended Data Fig. 2a-e; Extended Data Fig. 3a-e). For tNS1-3G2 and tNS1-4B8 complexes, the 2D class averages suggest that each NS1 tetramer could bind four copies of Fab fragment (Extended Data Fig. 3a,b). To improve the final map quality, the density of NS1 tetramer and Fab variable region (Fv) was refined, yielding two maps with global resolution at 2.8Å and 2.6Å, respectively. The densities of NS1 tetramer and four Fv could be well traced and modeled (Extended Data Fig. 3a,b). The RMSD of the dimeric NS1 between our cryoEM structure and the crystal structure (PDB: 5O6B) reported previously is 0.98Å (Brown et al., 2016), suggesting the coupling of antibodies disrupt the structure of the dimeric NS1. The N-terminal *β*-roll domain (residues 1–29), the wing domain (residues 30–180), the C-terminal *β*-ladder domain (residues 181–352), and the connector subdomain (residues 30-37 and 152-180) of NS1 could be well distinguished in our structures. The stacking of tetrameric NS1 is mediated by two dimeric NS1 at the inner hydrophobic surface comprised of central *β*-roll and extended the connector subdomain (Fig. 1a,b).

**Fig 2.**
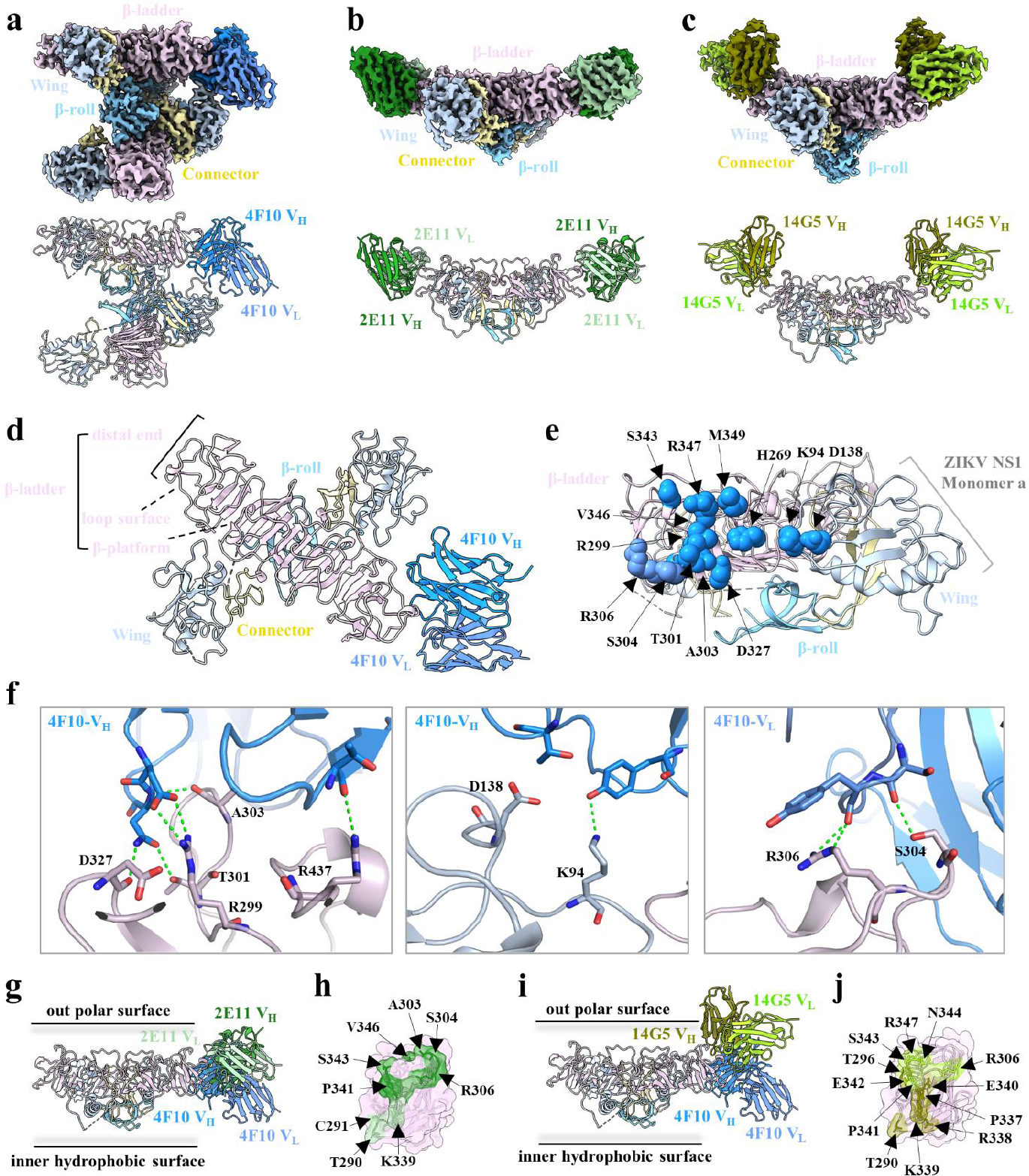
CryoEM structure analysis of ZIKV NS1 tetramer in complex with Fab 4F10, Fab 2E11 and Fab 14G5;s. (a-c) Overview of the density map and atomic model of ZIKV NS1 tetramer with two 4F10 Fv fragments, ZIKV NS1 dimer with two 2E11 Fv fragments, and ZIKV NS1 dimer with two 14G5 Fv fragments. The NS1 is colored as in Figure 1. 4F10 is colored dodger blue (4F10 V_H_) and cornflower blue (4F10 V_L_). 2E11 is colored forest green (2E11 V_H_) and dark sea green (2E11 V_L_). 14G5 is colored olive (14G5 V_H_) and yellow green (14G5 V_L_). This color scheme is used throughout the manuscript unless otherwise noted. (d) Zoomed-in view of the atomic model of the ZIKV NS1 dimer and 4F10 complex; In this structure, the left monomer is dark gray, and the right monomer is color-coded by domain as in panel a. (e) Mapping of critical residues for 4F10 on ZIKV NS1 dimer in top view. The dimer is color-coded as in panel c. Critical residues are indicated by spheres and are color-coded following the mAb panel in c. The epitopes of 4F10 are located on the distal end of the *β*-ladder domain, and wing domain of NS1 protomer. (f) Close-up view of the 4F10 epitope. Residues were shown as sticks; oxygen is colored red, nitrogen is colored blue, and hydrogen bonding is represented by dotted green line. Three panels from left to right are 4F10 V_H_ epitopes (left), 4F10 V_L_ epitopes(right); the distinct epitopes on NS1 wing domain occupied by 4F10 compared to 2E11 and 14G5 (middle). This panel was generated using PyMOL software. (g) Side-view of dimeric NS1 with 4F10 and 2E11. 2E11 does not create steric hindrance to the interface between the inner hydrophobic surface of the dimeric NS1 and the membrane. (g) Close-up view of the 2E11 epitopes on the distal end of the *β*-ladder domain. Side-view of dimeric NS1 with 4F10 and 14G5. 14G5 does not create steric hindrance to the interface between the inner hydrophobic surface of the dimeric NS1 and the membrane. (j) Close-up view of the 14G5 epitopes on the distal end of the *β*-ladder domain. Except for panel f, all panels were prepared using ChimeraX.

**Fig 3.**
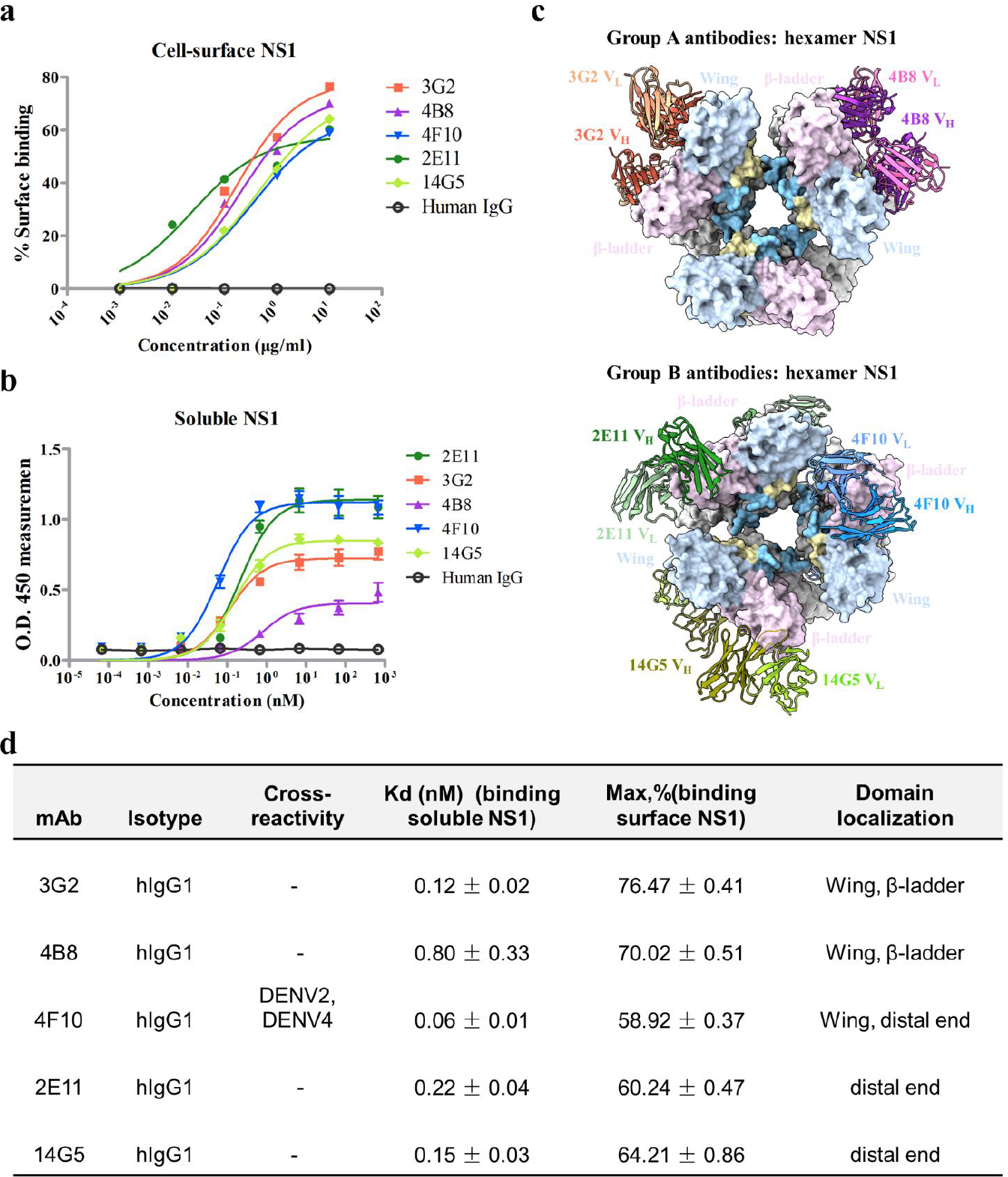
Binding affinity of 3G2 / 4B8 / 4F10 / 2E11 / 14G5 with the cell-surface and soluble ZIKV NS1 proteins; (a) The binding affinity of antibodies to cell-surface NS1 was determined by flow cytometry. The percentage of cell staining is shown as the average of three biological replicates in one experiment. Detailed methods and data are shown in Figure S8. (b) ELISA binding curves of 3G2 (orange), 4B8 (purple), 4F10 (blue), 2E11 (dark green), 14G5 (light green) to plate-bound ZIKV NS1. (c) Superimposition of group I and II antibodies onto the NS1 hexamer model (PDB: 4O6B). Antibodies and hexamer NS1 are shown in cartoon and surface representation respectively. (d) Summary of binding properties of anti-ZIKV NS1 mAbs.

The outer polar surface of dimeric NS1 is exposed to the solvent and occupied by two copies of Fab 3G2 or 4B8. As 3G2 and 4B8 bind at the outer surface of the tetrameric NS1 with nearly identical poses, we classified them into group I mAbs (Fig. 1a,b). As shown in Fig. 1c, one copy of 3G2 Fab occupies four regions on the outer surface of the dimeric NS1: the wing domain, the connector subdomain, and the middle of *β*-ladder domain of one monomeric NS1, and the middle of *β*-ladder domain of the other monomeric NS1. According to PISA calculations, the buried interface area between one 3G2 Fab and the dimeric NS1 is 1198Å^2^. Epitope residues of 3G2 are mapped onto the dimeric NS1 (Fig. 1d) and secondary structures (Extended Data Fig. 4). The variable loops of the 3G2 heavy chain (3G2 V_H_) contact with two regions: the connector subdomain (Asp174, Tyr175, Ser176, Leu177, and Glu178), and the *β*-ladder domain (Lys227, Trp232, Glu258, Glu315, and Thr317) (Fig. 1d,e). The hydrogen bonds coordinated by Tyr177, Cys178, and Glu315 stabilize the binding of 3G2 (Fig. 1e). The variable loops of the 3G2 light chain (3G2 V_L_) contact with Arg99, Pro101, and Gln102, and form a hydrogen bond to Lys146 on the wing domain (Fig. 1d,e). In addition, consistent with the results reported by Wesse et al., epitope residues Pro101, Lys146, Leu177, and Glu178 on the wing domain are important for protective mAbs to recognize NS1 on the membrane surface.

**Fig 4.**
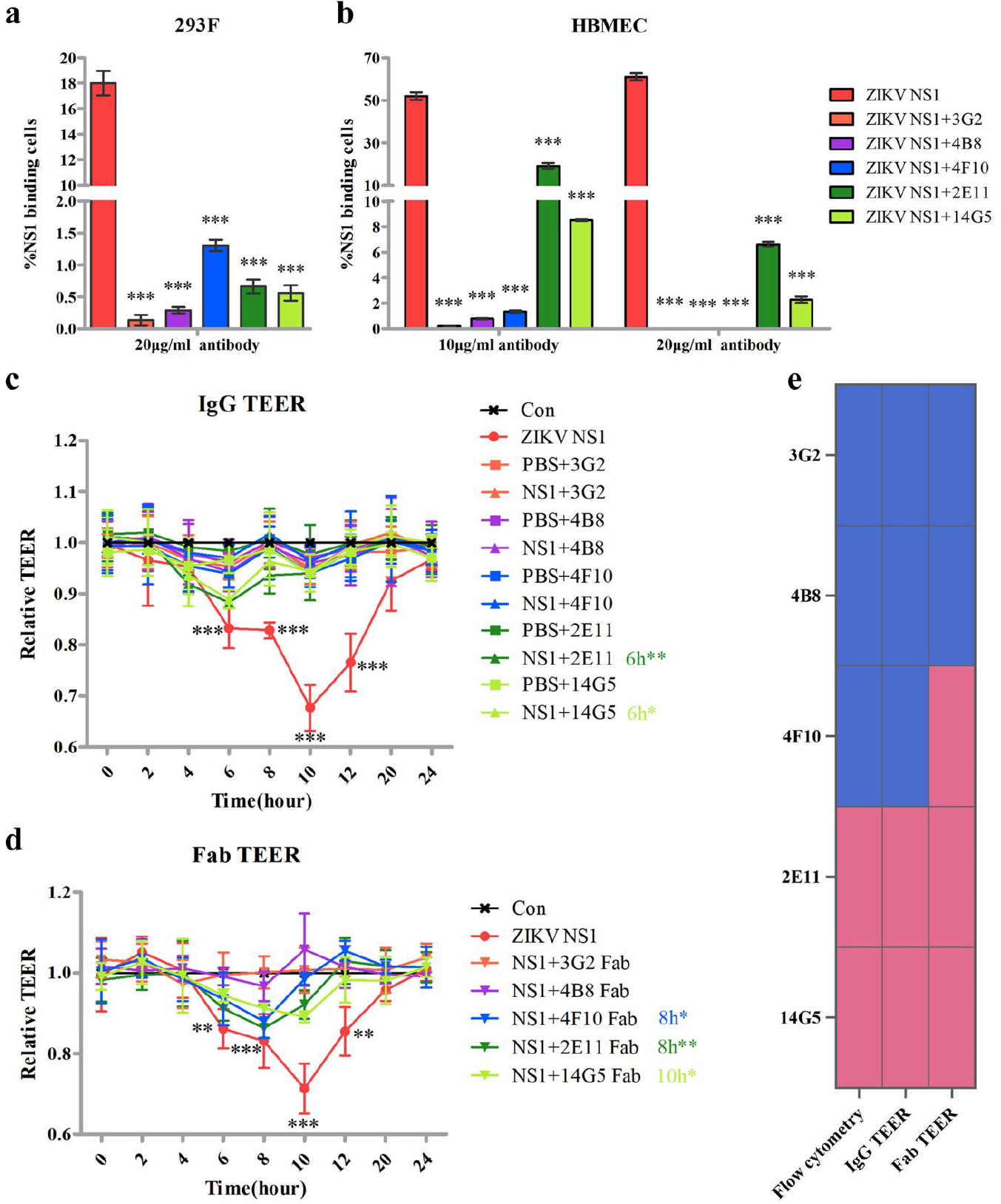
3G2 / 4B8 / 4F10 / 2E11 / 14G5 inhibit ZIKV NS1 cell binding and NS1-mediated endothelial hyperpermeability. (a, b) The cell binding experiments on 293F and HBMEC cell by pretreatment NS1 with five antibodies. 10,000 target cells were collected to analyze the proportion of FITC-positive cells. Data is shown as mean ± SD from three independent experiments. Using the Independent sample T-test to analyze the binding ability of antibodies, ***means P <0.001. (c, d) The IgG and Fab mAbs block NS1-mediated endothelial hyperpermeability in TEER assay.The concentrations of antibodies, Fab and native ZIKV NS1 were 20 ug/ml, 20 ug/ml and 5ug/ml, respectively. Data is shown as mean± SD from three independent experiments. The experimental group and control group were statistically significant by One-way ANOVA with Turkey’s multiple comparisons test, Black. “*” represents significant differences between the NS1 group and the Con group, Colored “*” with numbers represents significant differences between the antibody/Fab group and Con group at the time point, *means P<0.05, **means P<0.01, ***means p<0.001. (e) Schematic representation of neutralizing ability for Fab 3G2/4B8/4F10/2E11/14G5. Blue stands for high neutralization and pink stands for moderate neutralization. Decimal color codes in RGB are blue [73/108/206], pink [228/107/144].

The binding surface of 4B8 is similar to that of 3G2. The backbone of 4B8 V_H_ overlaps with 3G2 V_H_, while 4B8 V_L_ is slightly different from 3G2 V_L_ (Fig. 1f). The buried interface area between one 4B8 Fab and the dimeric NS1 is 979Å^2^, which is smaller than that of 3G2. As shown in Fig. 1g and Extended Data Fig. 4, and Extended Data Fig. 5, the majority of epitope residues recognized by 4B8 are consistent with 3G2. The interaction between Arg99 in the wing domain of NS1 and 3G2 V_H_ is not observed and the hydrogen bond with Glu315 on the *β*-ladder domain of NS1 is missing in the structure of the tNS1-4B8 complex (Extended Data Fig. 5).

In addition, sequence alignment showed that most epitope residues of group I mAbs are not conserved across flaviviruses (Extended Data Fig. 4). Further enzyme-linked immunosorbent assay (ELISA) analysis confirmed that 3G2 and 4B8 have no cross-reactivity with other flavivirus family members (Extended Data Fig. 6). These results suggest that group I antibodies have potential as diagnosis tools and their epitopes mode may be used for developing species specific antibodies for flaviviruses.

### Group II Antibodies (4F10, 2E11, and 14G5) Target Epitopes Residues at the Distal End of *β*-ladder Domain of NS1

We also determined the cryoEM structures of the NS1 combined with the Fab fragment of 4F10, 2E11, and 14G5. (Extended Data Fig. 3c-e). For the tNS1-4F10 complex, the models for the NS1 tetramer and two Fabs at the variable regions could be built while the densities of the other two Fabs are blurring (Fig. 2a and Extended Data Fig. 3c). For the dNS1-2E11 and dNS1-14G5 complex, the final 3D refinements were performed with a mask on one NS1 dimer and Fv of two Fabs due to the instability of tetrameric NS1 (Extended Data Fig. 3d,e). Therefore, the models of one NS1 dimer with two Fv were built for the dNS1-2E11 and dNS1-14G5 complex (Fig. 2b,c).

Both the dNS1-4F10 complex and the dNS1-2E11 complex have a straight bridge shape, whereas the dNS1-14G5 dimer complex presents an inverted arch shape. 4F10, 2E11, and 14G5 mainly recognize two loop regions (loop_290-306_ and loop_337-349_) in the distal end of the *β*-ladder domain (Fig. 2a-c), therefore they are categorized as the group II antibodies. The buried interface area between 4F10 Fab and the protomer NS1 is 798Å^2^. The epitopes at the distal end of the *β*-ladder domain play a major role in the binding of 4F10, and 4F10 forms additional interactions with the loop region of the wing domain (Fig. 2d). The variable loops of the 4F10 contact with residues Arg299, Ala303, Ser304, Gly305 and Arg306 in loop_290-306_, and residues Ser343, Val346, Arg347 and Met349 in loop_337-349_ of the *β*-ladder domain (Fig. 2e,f and Figure 2E, F, and Figure S7). The interactions between 4F10 V_H_ and residues (Lys94, Asp138, His269) in the wing domain from the same monomer NS1 were also observed in the structure (Fig. 2f, and Extended Data Fig. 7). It is worth noting that the majority of residues recognized by 4F10 are highly conserved in ZIKV and DENV2 (Extended Data Fig. 4). Epitope residues including Arg299, Ser304, Gly305, and Val346 have also reported in the recognition of broadly protective mAbs 1G5.3 and 2B7^24,25^. ELISA results have confirmed the cross-reactivity of 4F10 with DENV2 and DENV4 NS1 (Extended Data Fig. 6) ^27^, indicating 4F10 that is a potential broad-spectrum antibody.

Comparing the structures of 4F10, 2E11 and 14G5 have a translational shift towards the *β*-ladder domain and tilts towards the outer surface of the dimeric NS1 (Fig. 2g,i). The buried interface area of 2E11 Fab and 14G5 Fab with the protomer NS1 is 822.5Å^2^ and 772.6Å^2^, respectively. The interface between the wing domain and 4F10 is missing in the dNS1-2E11 and dNS1-14G5 complexes. The epitopes at *β*-ladder recognized by 2E11 include residues Thr290, Ala303, Ser304, and Arg306 in loop_290-306_, and Lys339, Pro341, Ser343, and Gln344 in loop_337-349_ (Fig. 2h and Extended Data Fig. 7b). The epitopes recognized by 14G5 are similar but with some variation from 2E11. The variable loops of the 14G5 interact with residues Thr290, Thr296, and Arg306 in loop_290-306_, and residues Pro337-Gln344 and Arg347 in loop_337-349_ (Fig. 2j and Extended Data Fig. 7c). Although 2E11 and 14G5 bind to the distal end of the *β*-ladder domain (loop_290-306_ and loop_337-349_), their epitope residues are less conserved. ELISA experiments confirmed that these two mAbs have no cross-reactivity with other members of the flavivirus family (Extended Data Fig.6).

Besides the main interfaces described above, 4F10 has an additional contact with a wing domain of the second dimeric NS1. The variable loops of the 4F10 interact with residues Tyr122, Val124, and Ala127 in the wing flexible loop and Glu80 in another peripheral loop in the wing domain. Besides the Glu1 in the variable loop in 4F10 forms a hydrogen bond with the residue Lys128 in wing flexible loop (Extended Data Fig. 7a). The structure of the tNS1-4F10 complex is more stable while the structures of tNS1-2E11 and tNS1-14G5 complex are highly flexible in the cryoEM analysis. The additional binding sites may contribute to stabilize the structure of tNS1-4F10 complex. Overall, our results showed that 4F10 can simultaneously interact with two dimeric NS1 from the tetrameric NS1, suggesting it may provide better efficiency in blocking the toxicity of NS1 than other group II antibodies.

### Binding properties of five anti-ZIKV NS1 mAbs with cell-surface NS1 and sNS1

NS1-targeting protective mAbs may function in two ways: clearing membrane-associated NS1 in ZIKV-infected cells through Fc receptor-mediated cellular immunity ^20,26^; or directly binding sNS1 to block it from entering the cell. ^21,24^. To understand the characterization of these five antibodies, we first examine the affinity of five mAbs with the membrane-associated NS1 by flow cytometry (Fig. 3a and Extended Data Fig. 8). The results showed that both group I and group II mAbs recognize the cell surface NS1 in a dose-dependent manner and the binding capacity of two group mAbs are maximized at a dose of 10 ug/ml (3G2: 76%; 4B8: 70%; 4F10: 58%; 2E11: 59%; 14G5: 63%). Although mAbs within the same group shared comparable capacity, group I mAbs showed stronger binding capacity with cell surface NS1 than group II mAbs (Fig. 3a and Fig. 3d). This suggests that the binding mode of group I mAbs is benefit to recognize the NS1 in cell surface.

Recent studies suggested that secreted NS1 may exist as oligomers including dimer, tetramer, and hexamer. Our structures show that both group I and group II Fabs bind to the dimeric NS1 and tetrameric NS1 at in a ratio of 1:1. To illustrate the interaction of the antibodies with hexamer, our structures were superimposed on a proposed model of NS1 hexamer (PDB: 4O6B). The docking results showed that all five Fabs interact with the hexameric NS1 without steric hindrance (Fig. 3b). Therefore, we suggest that both group I and group II antibodies can recognize epitopes in all of the oligomeric NS1. We further used ZIKV-derived sNS1 to evaluate the binding affinity of two groups mAbs by ELISA. The dissociation constant (Kd) values indicate that 4F10 binds to soluble NS1 with the highest affinity (Kd: 0.06 ± 0.01 nM), followed by 2E11 (Kd values of 0.22 ± 0.04 nM), 3G2 (Kd values of 0.12 ± 0.02 nM), 14G5 (Kd values of 0.15 ± 0.03 nM), and 4B8 (Kd values of 0.80 ± 0.33 nM). (Fig. 3c,d). Although all five mAbs retain nanomolar affinity for sNS1, 4F10 stands out with a 2-10 fold higher affinity than other mAbs. We speculate that the unique binding mode of 4F10 allow it to interact with the wing domains of two dimeric NS1 simultaneously (Extended Data Fig. 7a), contributing to an additional interface than other antibodies to block the soluble NS1.

### Group I and group II antibodies exhibit distinct protective mechanism in abrogating NS1-mediated endothelial dysfunction

To differentiate function among these antibodies in inhibiting cellular interaction of NS1 in vitro, we performed NS1 cell-binding experiments on 293F cells and human brain microvascular endothelial cells (HBMECs). Preincubating sNS1 with group I mAbs completely blocks ZIKV sNS1 binding to 293F cells and HBMECs regardless of high dose (20 *μ*g/mL) or low dose (10 *μ*g /mL) (Fig. 4a,b and Extended Data Fig. 9). Since group I antibodies bind to the outer surface of dimeric NS1(Fig. 1f), they are unlikely disrupt the interaction between the hydrophobic surface of dimeric NS1 and the cell membrane. We speculate that the binding mode of group I antibodies may represent a new protective mechanism. Epitopes of group I antibodies are widely distributed in the wing domain and *β*-ladder domain, indicating it may simultaneously disturb the interactions mediated two NS1 domains. It may block sNS1 from interacting with the endothelial cell surface, thereby preventing endothelial dysfunction and leakage.

For group II mAbs, preincubation of NS1 with 4F10, 2E11, and 14G5 significantly blocks NS1 binding to 293F cells (Fig. 4a). However, the dose-response analysis showed that only 4F10 were sufficient to completely abolish sNS1 toxic activity in HBMEC cells, while 2E11 and 14G5 partially inhibit NS1-mediated cell binding even at high dose (20 μg/mL) (Fig. 4b). The ability of these antibodies to inhibit NS1-mediated cell binding may vary depending on the different epitopes and affinity for sNS1. Both group II antibodies and the previously reported antibodies (2B7/1G5.3) bind to the distal end of *β*-ladder domain (Extended Data Fig. 10a,b), suggesting that they may share a similar blockade mechanism, such as inhibiting the conformational change of NS1 required for cell entry.

Protective antibodies against ZIKV NS1 confer protection through both Fc receptor-dependent and - independent pathways ^6,15,26^. To elucidate the relationship between the epitope recognition and differential protective mechanism, we the examined the protective effects of two groups antibodies (IgG and Fab) by measuring electrical resistance in a trans-endothelial electrical resistance (TEER) assay in HBMECs. Pretreatment of NS1 with both 3G2 and 4B8, either IgG or Fab, resulted in the complete elimination of NS1-induced endothelial hyperpermeability (Fig. 4d,e). The findings also validate the observation in vivo that destroying the Fc-mediated effector function of group I mAbs (3G2, 4B8) reduces but does not abolish their protective effects ^6^. For group II antibodies, 4F10 IgG mAb completely abrogated NS1-mediated cell barrier disruption in HPMECs (Fig. 4d,e), while 2E11 and 14G5 IgG mAb partially inhibit the function of sNS1 in HBMECs. Meanwhile, their Fabs were not sufficient to inhibit NS1-induced endothelial dysfunction. These findings indicate that the Fc regions of group II antibodies are required for completely abolishing NS1-mediated endothelial dysfunction. Compared to group II antibodies, the distinctive binding mode of group I antibodies presents potential benefits in damping NS1 toxicity in vitro regardless of IgG mAb or Fab.

## Discussion

Previously, NS1-specific mAbs (3G2, 4B8, 4F10, 2E11, and 14G5) were isolated from ZIKV convalescent patients and the protection of 3G2, 4B8, and 4F10 against ZIKV infection was evaluated in a neonatal mouse model^6,27^. Nonetheless, the differential protective mechanisms between group I mAbs (3G2 and 4B8) and group II mAb (4F10) are still unclear. In this study, we illustrated the epitopes of the five human anti-NS1 antibodies by solving the cryo-EM structures at atomic resolution. The purified sNS1 from S*f*9 cells was found to mainly exist as a tetramer, consistent with the finding that the recombinant DENV2 sNS1 exists predominantly in the tetrameric form from 293F cell^28^. Our data provided detail on how five antibodies recognize the epitopes of multiple oligomeric sNS1 and the validation the function of antibodies.

Our studies reveal a novel binding mode of group I mAbs and clarify how these antibodies provide excellent protection against ZIKV infection. Group I antibodies occupy approximately 2000Å^2^ interface area of the dimeric NS1 and recognize a large cluster of epitope residues on the outer surface. The avidity for cell surface NS1 for group I mAbs may contribute to induce antibody dependent cell-mediated cytotoxicity (ADCC) for clearing virus-infected cells. Notably, most of the epitope residues of group I mAbs are not conserved and ELISA analysis confirmed that group I mAbs have no cross-reactivity with other flaviviruses. Moreover, both IgG mAb and Fab of group I antibodies showed superior efficacy in abrogating NS1-triggered endothelial dysfunction. These results revealed a novel protective mechanism of group I antibodies, and it is worth exploring their potential as diagnostic tools and for the development of species-specific antibodies against flaviviruses.

For group II mAbs, some epitope residues were highly conserved in multiple flaviviruses. However, only 4F10 has cross-reaction with DENV2 and DENV4 NS1. The characteristics of group I and group II mAbs shows the possibility to combine these antibodies for develop complementary diagnostic tools in the serological discrimination of ZIKV. In addition, 4F10 draws our attention with its ability to block NS1-mediated membrane association. On the one hand, mAb 4F10 has the highest affinity with soluble NS1 so that it can provide better protection to abolish the NS1-mediated cell binding on HBMEC cells with the efficiency equivalent to group I antibodies (Fig.3b); On the other hand, 4F10 have extra contact with the wind domain of NS1 comparing to the other two antibodies (Fig. 2e,f), indicating 4F10 may create directly steric hindrance of NS1 to simultaneously inhibit the wing domain mediated cell binding and *β*-ladder triggered downstream events.

In summary, these data expand our understanding of the protective mechanisms of NS1-targeting antibodies with distinct binding modes and provide exact epitope information for the rational design of NS1-based antibodies and vaccine development.

